# Differential patterns of automatic segmentation of 3T and 7T MRI: Implications in neuropsychiatric disease

**DOI:** 10.1101/2023.01.25.525542

**Authors:** Gaurav Verma, Ki Sueng Choi, Helen Mayberg, Priti Balchandani

## Abstract

Automatic segmentation was performed on T1-MPRAGE structural MRI data acquired at 3T and 7T from 37 and 69 distinct healthy controls, respectively. Additionally, segmentation was performed on imaging acquired from 215 major depressive disorder (MDD) patients at 3T and 40 MDD patients at 7T. Of 259 segmentation-derived imaging features evaluated, 120 showed significant 3T vs. 7T differences among controls, and 153 among patients. 7T imaging metrics showed consistently lower cortical thickness and cortical gray/white matter ratios. Subcortical and cortical volumes measured at 7T were more mixed, with 7T images showing greater frontal lobe volume, but lower cortical volumes elsewhere.

## Introduction

Ultrahigh field MRI (≥7T field strength) facilitates imaging of brain structures with greater resolution and contrast than images acquired at more conventional field strengths of 1.5-3T. This improved detail enables surface-based morphometry with greater precision, potentially enabling segmentation of small brain structures such as thalamic nuclei and hippocampal subfields. Morphometry of these structures may yield imaging-derived biomarkers useful in the assessment of neuropsychiatric diseases such as major depressive disorder (MDD).

However, ultrahigh field MRI may present contrast levels different from 1.5-3T images, due both to B_1_-inhomogeneities and altered tissue contrasts. Since automated segmentation algorithms like FreeSurfer are optimized to ranges of timing parameters and tissue contrasts more typical of 1.5-3T, segmentation of ultrahigh field MRI may result in different segmentation patterns due to these unexpected properties. If these patterns systematically overestimate or underestimate morphological metrics, it may introduce systemic bias to the brain models derived from them. This study performed FreeSurfer automatic segmentation on cohorts of healthy controls and MDD patients at both 3T and 7T and assessed patterns of variation between those field strengths, both in individual metrics and whole-brain measures.

## Materials & Methods

T_1_-MPRAGE structural imaging was acquired from 69 healthy controls (24F/45M, mean age 36.5±10.5 years ranging from 20 to 56). Scan parameters included: Siemens 7T MAGNETOM, 32Rx/1Tx head coil, 1.95ms/3000ms TE/TR, 225×183mm field-of-view (FOV), 224 slices, 0.7mm^3^ isotropic resolution, 7 minute scan time. T_1_-MPRAGE was also obtained from a separate cohort of 37 healthy controls (22M/15F mean age 36.9±8.4 years) at 3T. Scan parameters at 3T included: 3.02ms/2600ms TE/TR, 1.0 mm^3^ isotropic resolution, 256×256mm FOV, 176 slices, 5 minute scan time, body transmit, 32-channel receive head coil). Additionally, 210 patients diagnosed with MDD (124F/86M mean age 39.7±10.7 years) were scanned at 3T and 40 separate patients scanned at 7T using identical T_1_-imaging (16F/24M, mean age 38.9±11.3 years) and the data subsequently segmented with FreeSurfer version 6.0. In total, 259 imaging metrics were segmented spanning cortical thickness, volume and gray/white matter ratio and subcortical volume. Sixteen whole-brain metrics were considered, along with 68 cortical thickness metrics, 68 cortical gray/white ratios, 68 cortical volumes and 68 subcortical volumes. Although 7T images were acquired at higher native resolution, segmentation at both field strengths were performed without the high resolution flag activated, resulting in a final segmented resolution of 1 mm^3^ isotropic. The metrics were compiled and statistically analyzed in Matlab, and then visualized using the BrainNet library for Matlab.

## Results

Of 259 FreeSurfer-derived imaging features considered, 120 were significantly different (p<0.05) in comparisons between controls scanned at 3T vs. 7T. In comparisons among MDD patients, 153 of 259 imaging features showed significant differences between 3T vs. 7T. Field strength related differences in imaging features were largely the same whether in controls or MDD patients. Table 1 shows a summary of differences observed between 3T and 7T metrics and amongst patients and controls. Segmentation at 7T tended to consistently underestimate cortical gray/white ratio, and thickness, while differences in cortical and subcortical volumes were more mixed. Both amygdala and hippocampus showed patterns of underestimation at 7T among patients and controls. There were no instances in which a metric deviated in one direction among the control cohort and the opposite direction among patients.

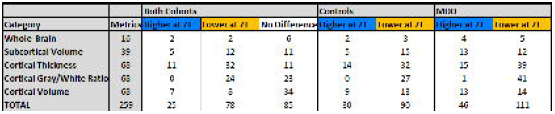

Figures 1 and 2 show heat maps of cortical regions indicating significant differences in one or more of three imaging metrics. Positive values, represented in blue, indicate significantly higher cortical volume, cortical thickness and/or cortical gray/white matter ratios at 7T compared to 3T. Negative values, represented in orange, show significantly lower values in one or more of those three metrics at 7T vs. 3T. Figures 3 and 4 show analogous metrics showing differences in subcortical volumes. Here, blue represents significantly higher subcortical volume at 7T and orange represents significantly lower subcortical volumes at 7T.

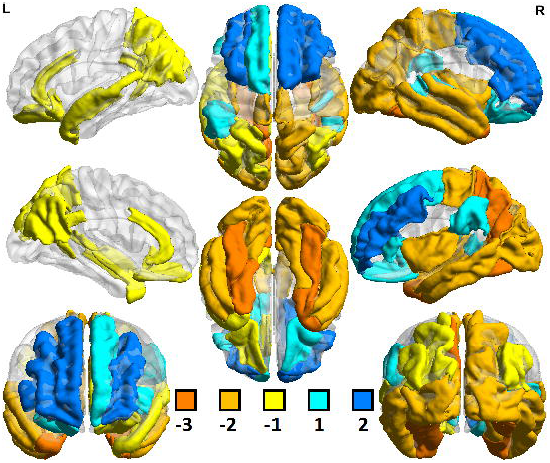

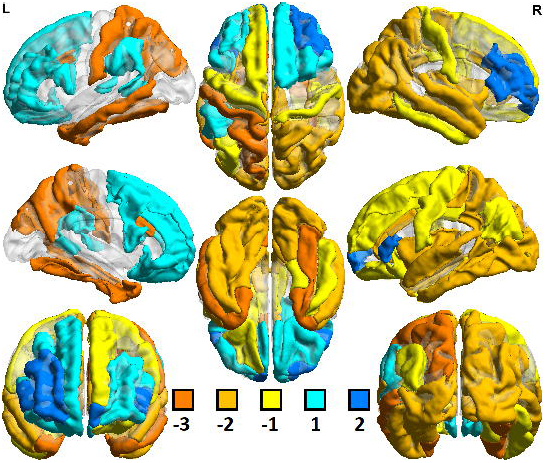

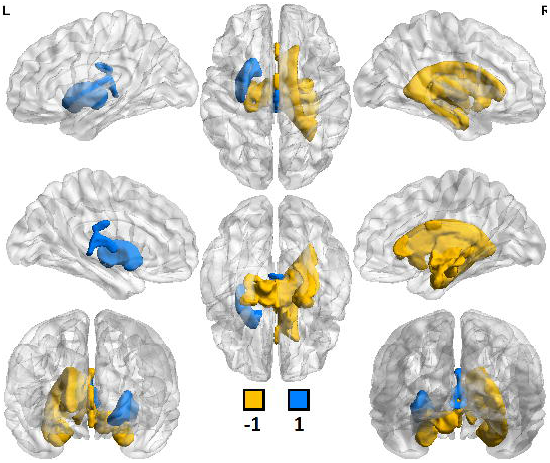

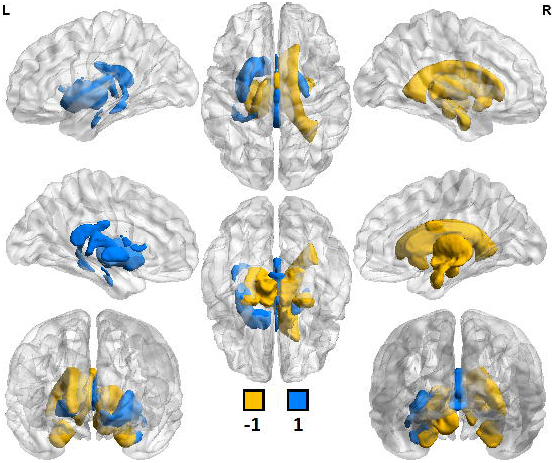

Global brain volume metrics like BrainSegVol, SupraTentorialVol and eTIV (representing total brain segmentation volume, supratentorial volume and total intracranial volume, respectively) showed no significant difference at 3T vs. 7T amongst both patients and controls. Instead, segmentation patterns showed consistently larger measures for cortical volume, thickness and/or gray/white ratio in the frontal lobe at 7T while the same metrics tended to be lower in the parietal, temporal and occipital lobes.

## Discussion

Ultrahigh field MRI enables acquisition of structural imaging with higher signal intensity and potentially higher resolution than scans acquired at 3T. However, subtle variances introduced by scanning at ultrahigh field may complicate segmentation of these images. B_1_ field inhomogeneities and the relatively faster tissue T_2_* relaxation time may result in segmented metrics with systematic differences dependent on field strengths. Patients and controls tend to show the same variations between the field strengths, suggesting these factors are not dependent on patient status. Therefore, comparisons between these cohorts are likely to be less subject to bias if scanned with the same field strengths. Previous research has also suggested imaging metrics like cortical thickness are systematically underestimated at 7T and that these underestimations can be more severe if the 7T imaging employs a higher native resolution than comparable 3T imaging, even if the segmented resolutions are the same. [1-5].

## Acknowledgements

The authors would like to acknowledge funding from NIH R01 MH109544 and the NARSAD Young Investigator Grant.

